# ANME-1 archaea drive methane accumulation and removal in estuarine sediments

**DOI:** 10.1101/2020.02.24.963215

**Authors:** Richard Kevorkian, Sean Callahan, Rachel Winstead, Karen G. Lloyd

## Abstract

Uncultured members of the Methanomicrobia called ANME-1 perform the anaerobic oxidation of methane (AOM) through a process that uses much of the methanogenic pathway. It is unknown whether ANME-1 obligately perform AOM, or whether some of them can perform methanogenesis when methanogenesis is exergonic. Most marine sediments lack advective transport of methane, so AOM occurs in the sulfate methane transition zone (SMTZ) where sulfate-reducing bacteria consume hydrogen produced by fermenters, making hydrogenotrophic methanogenesis exergonic in the reverse direction. When sulfate is depleted deeper in the sediments, hydrogen accumulates making hydrogenotrophic methanogenesis exergonic, and methane accumulates in the methane zone (MZ). In White Oak River estuarine sediments, we found that ANME-1 comprised 99.5% of 16S rRNA genes from amplicons and 100% of 16S rRNA genes from metagenomes of the Methanomicrobia in the SMTZ and 99.9% and 98.3%, respectively, in the MZ. Each of the 16 ANME-1 OTUs (97% similarity) had peaks in the SMTZ that coincided with peaks of putative sulfate-reducing bacteria *Desulfatiglans sp.* and SEEP-SRB1. In the MZ, ANME-1, but no putative sulfate-reducing bacteria or cultured methanogens, increased with depth. Using publicly available data, we found that ANME-1 was the only group expressing methanogenic genes during both net AOM and net methanogenesis in an enrichment. The commonly-held belief that ANME-1 perform AOM is based on the fact that they dominate natural settings and enrichments where net AOM is measured. We found that ANME-1 also dominate natural settings and enrichment where net methanogenesis is measured, so we conclude that ANME-1 perform methane production. Alternating between AOM and methanogenesis, either in a single ANME-1 cell or between different subclades with similar 16S rRNA sequences of ANME-1, may confer a competitive advantage, explaining the predominance of low-energy adapted ANME-1 in methanogenic sediments worldwide.

**Abstract Importance:** Life may operate differently at very low energy levels. Natural populations of microbes that make methane survive on some of the lowest energy yields of all life. From all available data, we infer that these microbes alternate between methane production and oxidation, depending on which process is energy-yielding in the environment. This means that much of the methane produced naturally in marine sediments occurs through an organism that is also capable of destroying it under different circumstances.

## Main Text

Non-seep marine sediments are the third largest producers of methane on Earth, after rice production and wetlands (1). However, very little of this methane reaches the ocean floor because abundant uncultured archaea of the Methanomicrobia group, called ANaerobic MEthane oxidizers (ANME), catalyze the anaerobic oxidation of methane (AOM) (2, 3). However, the identities of the organisms that produce the majority of this methane are largely unknown in non-seep sediments, since cultured methanogenic groups are often difficult to find in methanogenic depths of coastal sediments (4). Some researchers have suggested that ANME can also be capable of methane production, mostly from studies showing that they are the most abundant putative methane-metabolizing organisms in methane-producing non-seep sediments (5–9). However, sampling with a fine depth resolution as sediments switch from net methane removal to net methane production is required to determine whether ANME-1 might drive this shift. This is because the total population sizes of ANME-1 or other likely methanogens may be dynamic over spatial scales that are missed by studies examining only a few depths distributed throughout a sediment column.

Most microbes in marine sediments belong to uncultured genera to phyla (10), making it necessary to infer their physiologies in a natural setting rather than in axenic cultures. Some of these microbes drive sulfate reduction and methanogenesis, which are the key respiratory processes that directly or indirectly oxidize organic matter in anoxic marine sediments. The balance between diffusion and biological respiration drives a down-core shift from sulfate reduction to methanogenesis, with net removal of methane through AOM in the sulfate methane transition zone (SMTZ) at intermediate depths (11). Methanogens conserve energy by producing methane from hydrogen plus carbon dioxide, acetate, formate, or a range of methylated compounds. When methanogenesis is produced from hydrogen plus carbon dioxide, it is referred to as hydrogenotrophic methanogenesis (Eq. 1). In the SMTZ, sulfate reducers keep hydrogen concentrations low enough to make hydrogenotrophic methanogenesis exergonic in the reverse direction (12). This is possible because hydrogenotrophic methanogenesis, unique among many other respiratory mechanisms, can be made to be exergonic in the reverse direction by changing the relative concentrations of products and reactants. Hydrogenotrophic methanogenesis (Eq. 1) is reversible because 1) it has a stoichiometry of 4 molecules of hydrogen per methane molecule, giving hydrogen activities (activity is similar to concentration, after accounting for the ionic strength of the solution) a large amount of power over the thermodynamics and 2) hydrogen activities in marine sediments can be extremely low, down to ~10^−9^ times the standard state activity (13). *In situ* measurements of porewater chemistry have demonstrated this reversal of ΔG between the SMTZ and the methane-containing zone when hydrogen concentrations decrease (14).

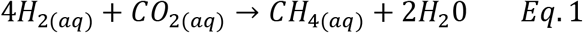

While some metabolic pathways such as the citric acid cycle are amphibolic (15), meaning they can occur in either the catabolic or anabolic direction, it is unknown whether a single organism can conserve energy from either hydrogenotrophic methanogenesis or AOM, depending on which direction is exergonic. Although cultured methanogens have been shown to catalyze methane oxidation, they cannot sustain the process, leading Valentine et al. to conclude that AOM does not occur through a reversal of the methanogenic metabolic pathway in most of the commonly-studied methanogenic strains in culture (16). However, it is possible that these methanogens could not sustain hydrogen production because they were adapted to very high energy yields (17) and could not survive on the paucity of energy afforded by AOM. ANME-1 archaea, on the other hand, have low energy requirements (18), making them good candidates for being reversible methanogens. ANME-1 are present and active in both AOM and methanogenic zones (8, 19–21), and they are phylogenetically related to cultured methanogens, belonging to the Methanomicrobia, a group for which all cultured strains are methanogens (22). Initial incomplete genomes from ANME-1 contained all of the genes required for methane production except for the one encoding N5, N10-methylene-tetrahydromethanopterin reductase (*mer*) (23, 24). More recently, *mer* has been found in ANME-1 genomes (20). ANME-1 has been shown to be enriched during methane production (5), and its biomass has variable stable carbon isotope values in nature, suggesting that it can use substrates other than methane to make biomass (7).

In order to infer physiology of uncultured ANME-1, we examined its population growth dynamics in fine-scale depth resolution in sediments of the White Oak River estuary and laboratory enrichments with publically available data (25). Sediments experiencing a consistent and known sedimentation rate offer an opportunity to substitute depth for time and test for the increase of a microbial population over a particular depth interval. An increase in total population size with depth implies that net growth must have occurred, although it may have been faster than the measured increase rate due to an unknown death rate. Similarly, a decrease in population size with increasing depth implies net death. Quantifying the absolute abundance of a particular microbial population in marine sediments, however, is inaccurate with current methods such as 16S rRNA gene amplicon libraries, quantitative PCR (qPCR) and Fluorescent In Situ Hybridization (FISH) (26–28). However, two of these methods, 16S rRNA gene amplicon libraries and qPCR, are quantitative in relative terms when comparing samples with similar DNA extraction and amplification biases (27, 28). Therefore, in order to measure increases or decreases in population size with depth, one can determine the change in the Fraction Read Abundance times total Cell count (FRAxC) with depth (29).

We examined 16S rRNA gene sequence FRAxC with high depth resolution (3 cm intervals in the upper 10 cm, 1 cm intervals between 10 and 60 cm, and then 3 cm intervals from 60 to 80 cm below the sediment-water interface) throughout a diagenetic sequence in sediments of the marine-influenced White Oak River estuary. Due to sediment volume restrictions from the small depth intervals, 16S rRNA gene amplicon libraries, cell counts, and hydrogen concentrations were measured for one core (core 6), sulfate concentrations were measured for the other (core 1), and methane concentrations and δ^13^C values were measured on both cores. The fine depth resolution allowed us to visualize any die-off and regrowth of ANME-1 between AOM and methanogenic zones, which would suggest that different ANME-1 populations perform each of the two metabolisms. The finescale approach also allowed the potential detection of cultured methanogens in any of the sediment layers that may have been missed in previous studies of estuarine sediments.

## Results/Discussion

Total cell abundance decreased sharply within the first 10 cm from 2.0 × 10^8^ to 2.6 × 10^7^ cells/g and remained steady at ~1.3 × 10^7^ cells/g for the rest of the core (Fig. 1a). Sulfate concentrations in core 1 decreased from a surficial concentration of 11.2 mM to a constant concentration of 0.06 ± 0.02 mM at 65 to78 cm (Fig. 1e). This concave decrease in sulfate is consistent with it being consumed by sulfate-reducing bacteria (30). Aqueous methane concentrations were ~0.004 mM in near-surface sediment and increased to >0.6 mM at 60 cm in core 6 (Fig. 1f) and 75 cm in core 1 (Fig. 1b), with a generally concave shape indicating AOM. Below this, methane increased to 0.73-0.87 mM in the 75-78 cm depth interval of both cores. In core 6, methane concentrations the methane concavity shifted between ~55 cm and 73 cm, consistent with methanogenesis. Between 2.5 cm and 64.5 cm, hydrogen concentrations were low (0.07 - 2.05 nM) in core 6 (Fig. 1d), consistent with those predicted for sulfate reducers operating at their minimum energy (1.22 ± 0.45 nM in similar sediments, which yields a ΔG of - 20 kJ/mol sulfate, just slightly above the minimum free energy conservation (13)). Below this, hydrogen increased above 6 nM, higher than the threshold for energy conservation for hydrogenotrophic methanogenesis at 5.11 nM in similar sediments (13). In core 1, δ^13^C of methane decreased from −41‰ ± 1.17 at 41.5 cm to −72‰ ± 0.10 at 73.5 cm (Fig 1g). In core 6, δ^13^C of methane decreased from −46‰ ± 0.20 at 29.5 cm to −74‰ ± 0.02 at 67.5 cm (Fig. 1c). These values are consistent with the biogenic production of methane deeper in the sediments, resulting in enrichment of the lighter carbon isotope, ^12^C (31). Within the SMTZ, methane diffusing up through the sediment became gradually depleted in the lighter carbon isotope, consistent with AOM (32). This is because biological AOM has a preference for ^12^C over ^13^C, leaving the residual methane ^13^C-enriched (33, 34). The shift in δ^13^C-CH_4_ toward “heavier” values between ~60 and 30 cm indicates methane oxidation occurs in this depth interval. Methane concentrations in samples shallower than the AOM zone were too low to get accurate δ^13^C values, so the values in the upper parts of the cores were not used. Together, these geochemical measurements suggest the location of AOM (30 to ~60 cm) and methanogenesis (>60 cm) in core 6, the core where DNA measurements were made. The depth of the onset of net methanogenesis was chosen at ~60 cm because it is the depth where 1) sulfate is depleted in core 1, 2) hydrogen concentrations begin to continuously increase in core 6, 3) methane concentrations are concave-down, and 4) δ^13^C values become consistent with methanogenesis. The upper limit of AOM is defined by the depth at which methane concentrations begin to continuously increase downcore, which is 25 cm in core 6 and 45 cm in core 1. For consistency in nomenclature based on the available substrates, we refer to the depths where methane is present above 60cm the SMTZ, and the depths below 60cm the MZ, for methane zone.

**Figure 1.**
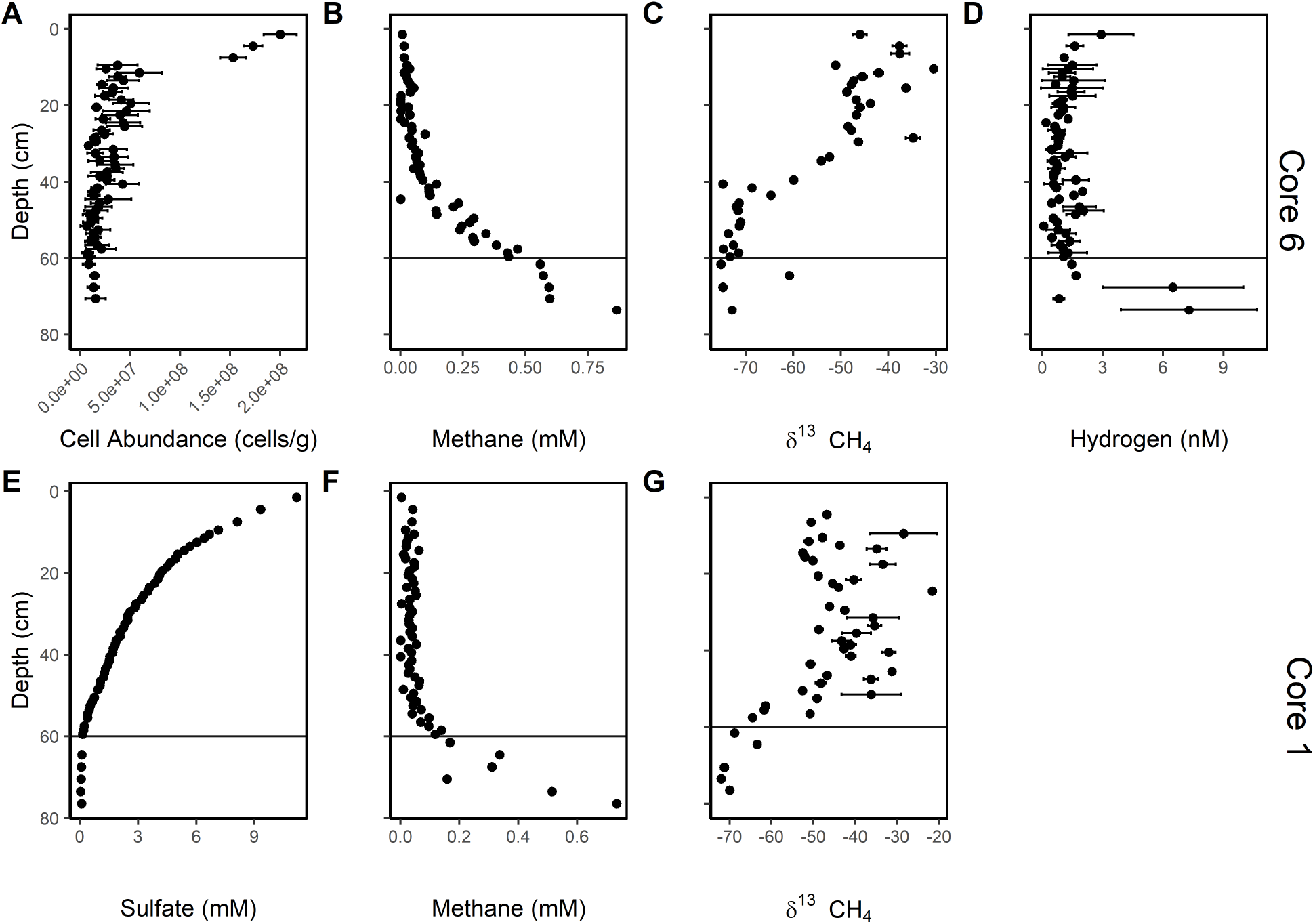
White Oak River estuary cores show methane-consuming or AOM sediments in the SMTZ and methane-producing or methanogenic sediments below it. Aqueous geochemistry for cores 6 (top row) and 1 (bottom row), with A) cell abundance, B and F) methane concentration, C and G) δ^13^C of methane, D) hydrogen concentration, and E) sulfate concentration. Black line shows approximate transition from AOM to methanogenesis.

ANME-1 dominated the Methanomicrobia in 16S rRNA gene amplicon libraries in both the SMTZ and MZ (99.5 and 99.9%, respectively, Fig. 2). In previous work, ANME-1 were shown to express methyl coenzyme M reductase subunit A (*mcrA*) genes, a key gene in methanogenesis and AOM (36), in both of these zones as well (8). ANME-1 comprised six OTUs of ANME-1a, nine OTUs of ANME-1b, and one ANME-1 OTU that could not be placed into one of those two subgroups. This agrees with previous observations from Aarhus Bay (20), White Oak River estuary (8), Gulf of Mexico deep-sea (19), and Santa Barbara Basin deep-sea (21) that ANME-1 were the dominant or only organisms with the genes for methane-cycling in AOM and methanogenic sediments. Each of these analyses utilized 16S rRNA primers capable of amplifying cultured methanogens, so the absence of cultured methanogens was not likely to be an artifact of primer bias. However, primer bias can greatly skew the relative abundance of different clades (26), so we analyzed 16S rRNA gene sequences from unamplified metagenomes from the White Oak River estuary (37) and found that ANME-1 comprised 100% of the Methanomicrobia in the AOM zone and 92.86% in the MZ, in agreement with amplicon data (Fig. 2).

**Figure 2.**
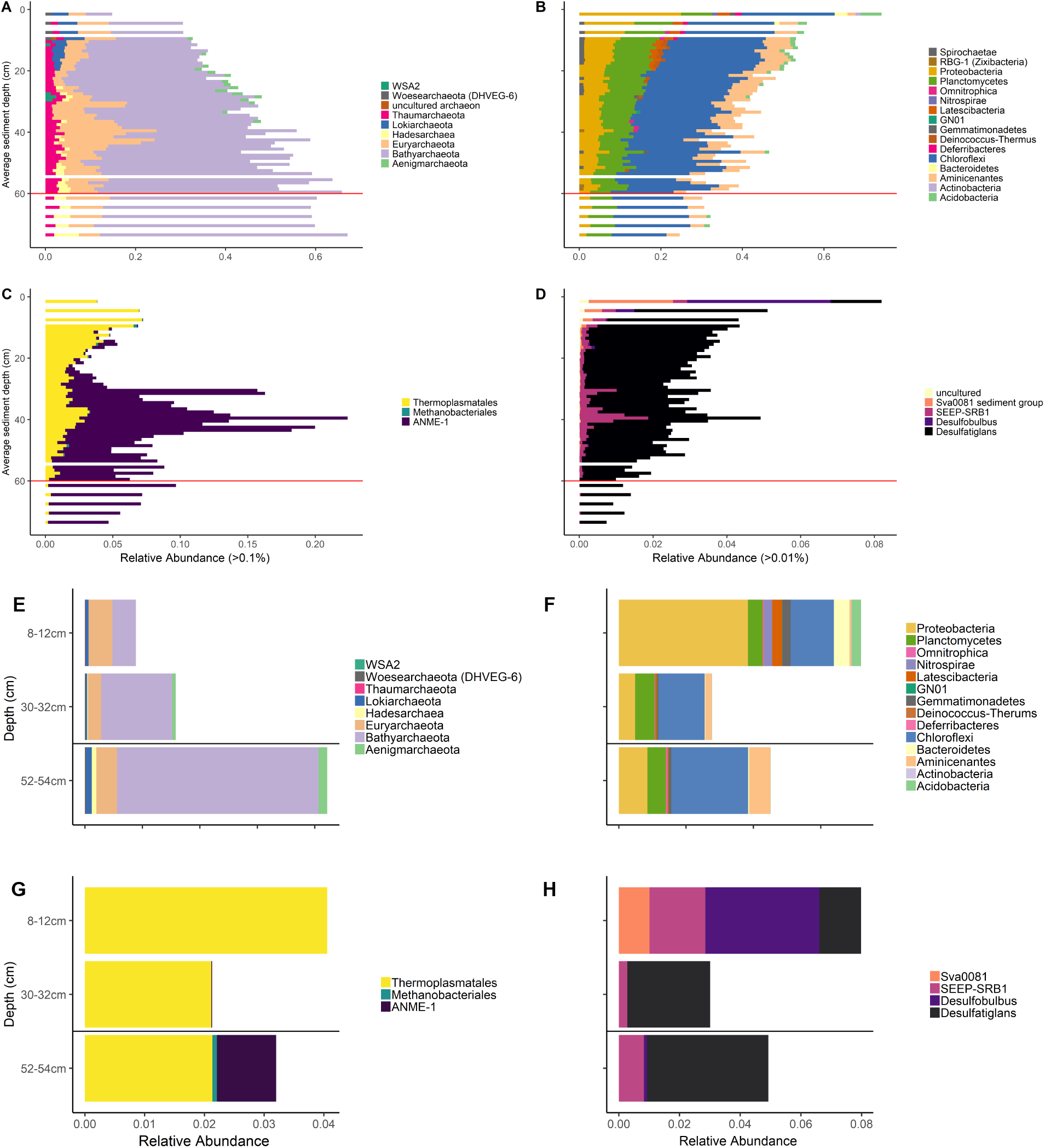
ANME-1 and *Desulfatiglans sp*. dominate methane- and sulfur-cycling organisms in both methane-consuming and methane-producing sediments. Relative abundance of 16S rRNA gene sequences for all archaea (left panels) and all bacteria (right panels), for amplicon libraries (A-D) and metagenomes (E-H), grouped at the phylum level. C, D, G, and H show putative sulfate-reducing bacteria and Euryarchaeota, grouped at the family level. Only phyla with >1% relative 16S rRNA gene sequence abundance for bacteria and > 0.1% for archaea are shown. Black lines show transition from AOM to methanogenesis.

The four most abundant ANME-1 OTUs, comprising 96% of all ANME-1 16S rRNA gene sequences, had one or more FRAxC peaks in the SMTZ that coincided with peaks in putative sulfate-reducing bacteria *Desulfatiglans sp.* and SEEP-SRB1 (Fig. 3). SEEP-SRB1 have been shown to form syntrophies with ANME-1 (25). *Desulfatiglans sp.* have not previously been associated with syntrophies with ANME-1, but were the most abundant sulfate-reducing bacteria in ANME-1-dominated methane seeps in the Gulf of Mexico (38). In White Oak River estuary sediments, *Desulfatiglans sp.* and SEEP-SRB1 comprised 99.9% of 16S rRNA gene sequences of known sulfate-reducing bacteria clades in both the SMTZ and MZ. They were distinct from the dominant clades present in the upper, methane-free sulfate reduction zone: *Desulfobulbus sp.* and the uncultured Sva0081 clade within the Desulfobacteraceae (94% of sulfate-reducing bacteria above 25 cm, Fig. 2). The dominance patterns of these potentially sulfate-reducing clades is supported by metagenomic data as well (Fig. 2). This suggests that *Desulfatiglans sp.* and SEEP-SRB1 may be adapted to syntrophy with ANME-1 in the SMTZ, and this may allow them to out-compete other sulfate-reducing bacteria deeper in the core when ANME-1 are abundant.

**Figure 3.**
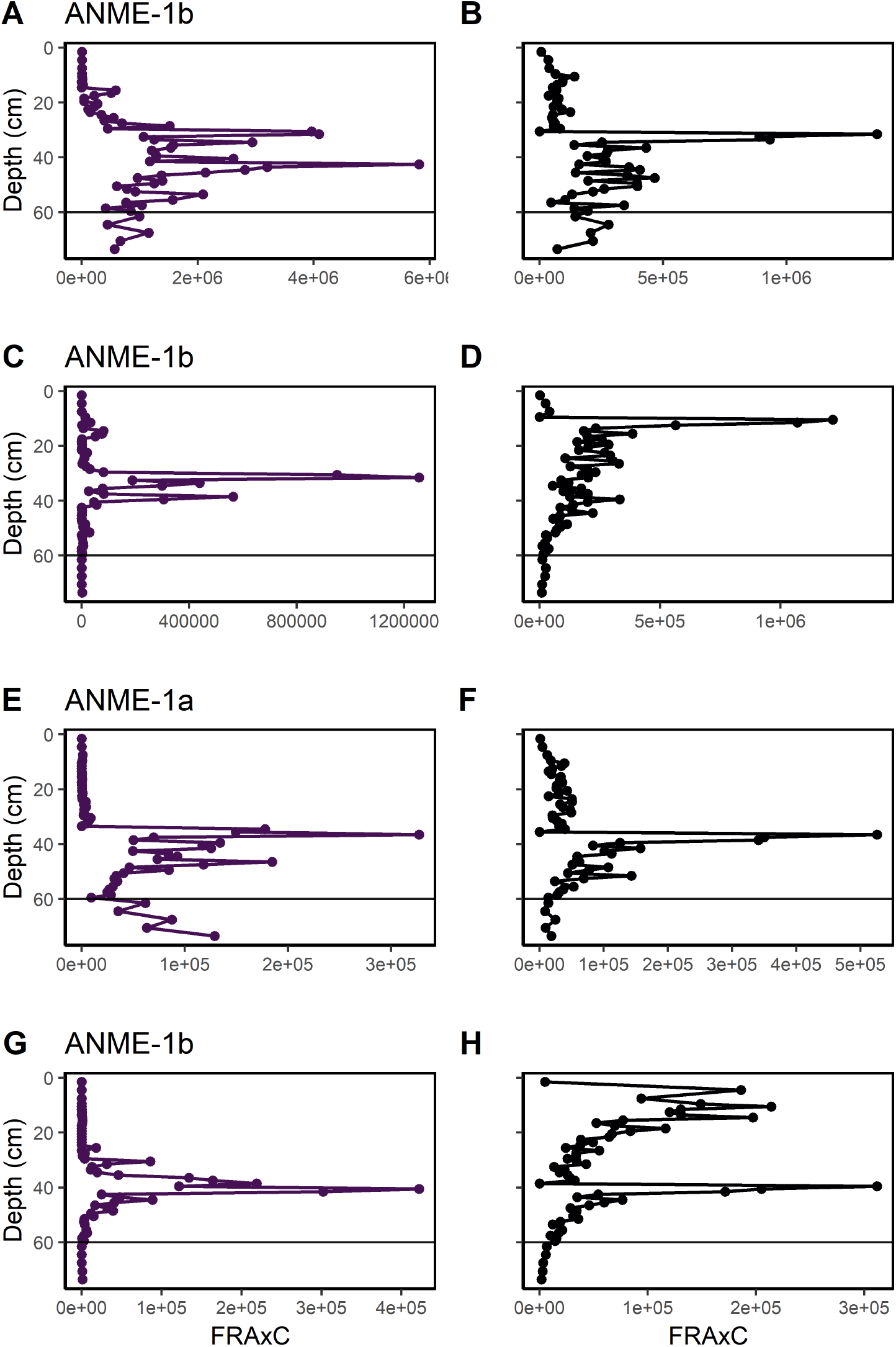
ANME-1 Fraction Read Abundance times Cell counts (FRAxC) and sulfate reducing bacteria have co-occurring peaks during AOM but not methanogenesis. Values for the four most abundant OTUs of ANME-1 (left panels, A, C, E, and G) and *Desulfatiglans sp.* (right panels, B, D, F, and H) with depth, ordered by decreasing OTU abundance from top to bottom panels. Colors match those in Figure 2, and black line indicates AOM to methanogenesis transition.

The most abundant ANME-1b OTU, which accounted for 83% of total ANME-1 reads, did not undergo a population decrease at the base of the SMTZ followed by an increase in the MZ (Fig. 3a), as was suggested in a previous experiment with lower depth resolution (8). This suggests that, for most ANME-1b, if they switch from methane-oxidizing to methane-producing, this either does not require a die-off followed by a separate population growing up, or the population decrease happens in a smaller depth interval than we could observe with 1 cm intervals. A third possibility is that a population decrease does occur between the SMTZ and MZ, but seasonal variability of SMTZ depth (8) dampens the coherence of this signal. Two ANME-1b OTUs decreased with depth in the MZ (Fig. 3b and 3d). This suggests that either reversibility is not universal among ANME-1 subpopulations, or that some subpopulations were outcompeted by other ANME-1 in the MZ. One ANME-1a OTU, which accounted for 3% of the total ANME-1 reads, did decrease at the base of the SMTZ and increase in the MZ, suggesting that die-off and regrowth may be required for some ANME-1 populations to switch between methane oxidation and production. In total, ANME-1 populations increased relative to those of sulfate-reducing bacteria throughout the transition from SMTZ to MZ (Fig. 4).

**Figure 4.**
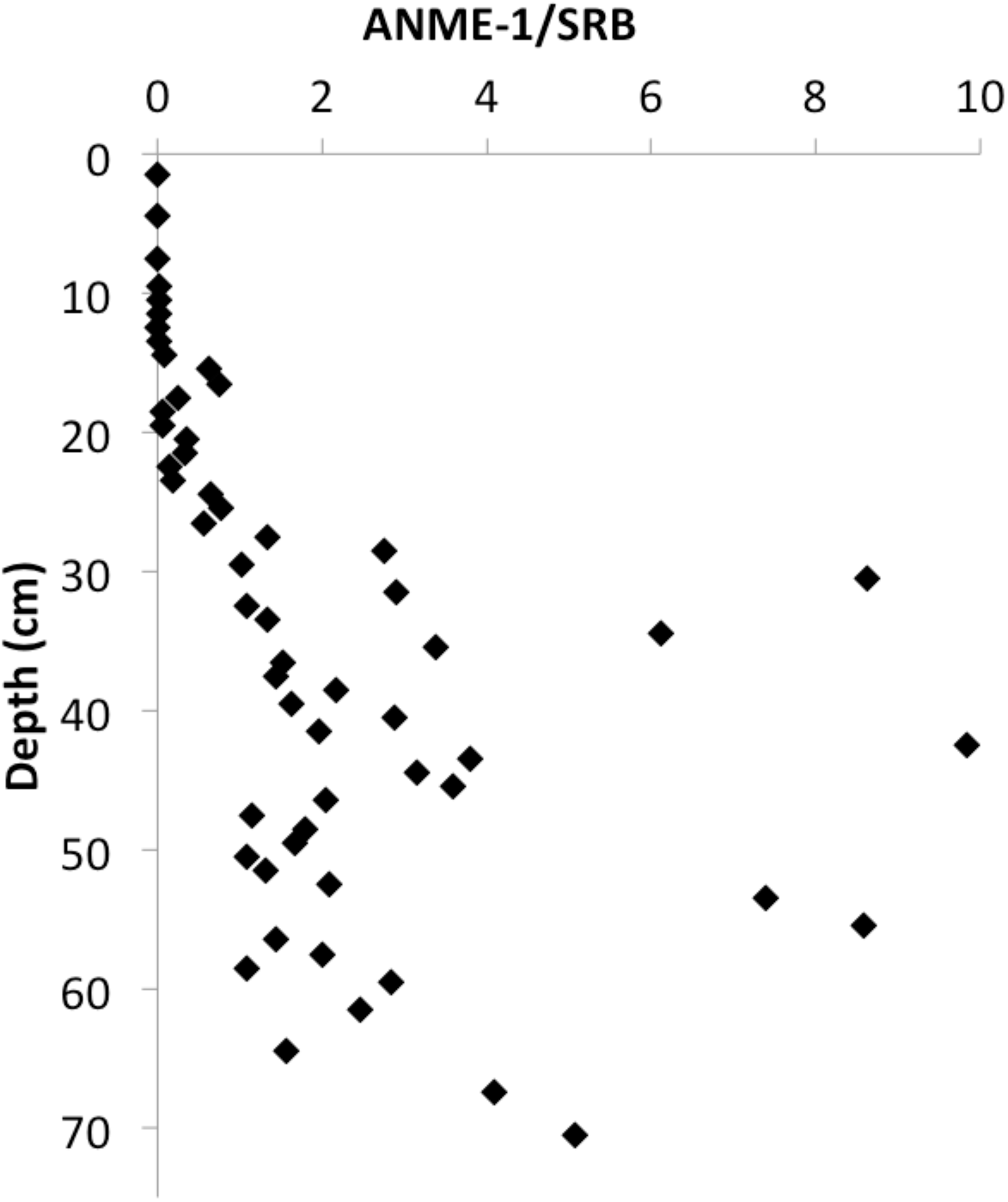
ANME-1 increased relative to sulfate reducing bacteria throughout the SMTZ and below. Ratio of total ANME-1 16S rRNA gene abundance to total sulfate reducing bacteria 16S rRNA gene abundance with depth.

The most abundant *Desulfatiglans sp.* OTU maintained its population through the MZ, and all others decreased, suggesting that successively smaller populations were capable of meeting their energetic needs on either cryptic sulfur cycling or fermentation as substrates were depleted with depth (Fig. 4). The coupling of ANME-1 and sulfate-reducing bacteria populations in the SMTZ, and their decoupling in the MZ, is consistent with ANME-1 switching from AOM, which requires a sulfate-reducing partner, to methanogenesis, which does not. ANME-1a and ANME-1b were three- and five-fold higher in the SMTZ than in the MZ. The amount of methane that is produced over many tens of centimeters of sediment depth in the MZ is consumed over just a few centimeters sediment depth in the AOM zone. Therefore, a reversible methanogen operating at similar cell specific metabolic rates in the SMTZ and MZ would be expected to be in a higher cell density in the SMTZ than in the MZ, as was observed.

The only non-ANME-1 member of the *Methanomicrobia* detected was one OTU of *Methanobacterales*, which ranged from 0 to 0.0026 % relative sequence abundance and decreased with depth in the MZ (Fig. 2), suggesting that it was not driving methanogenesis. The uncultured phylum, *Bathyarchaeota*, has been suggested to perform methane-cycling (39, 40) and increased in relative abundance with depth in the MZ (Fig. 2). However, the inference that *Bathyarchaeota* perform methane-cycling is based solely on the presence of an evolutionarily-divergent *mcrA* gene found in genomes from a terrestrial coal bed (39). Orthologs to this *mcrA* have been shown to catalyze butane rather than methane oxidation (41) and none of the *Bathyarchaeota* genomes obtained from the White Oak River estuary sediments have this gene (37). Instead *Bathyarchaeota* in marine sediments appear to perform acetogenesis and fermentation of organic substrates such as proteins and lignin (42, 43). Ten *Hadesarchaeota* OTUs increased in relative abundance in the MZ (Fig 2). This uncultured archaeal phylum has numerous carbon metabolism genes in common with *Methanomicrobia* but does not have a methanogenic enzymatic pathway. Instead *Hadesarchaeota* are hypothesized to have a heterotrophic and/or nitrogen cycling lifestyle (44). A proposed methyl-reducing methanogenic archaeal lineage, WSA2 (45), was also detected in our samples. However, the 11 OTUs were in very low relative abundance and declined below 27-30 cm (Fig. 2).

No bacteria or other archaea showed changes in relative or FRAxC abundance indicative of participation in methane cycling. OTUs from sulfide-oxidizing bacteria such as *Sulfurimonas*, *Thiotrichales*, and *Thiomicrospira* were either low in abundance, not detected, or demonstrated no significant increases in relative abundance with depth. Thirty-one OTUs of the organoheterotrophic genus *Caldithrix* and phylum *Defferibacteres* declined with depth (46). Obligate iron- and manganese-reducing bacteria either did not meet abundance thresholds or were not detected. Lokiarchaeota and Woesearchaeota both decreased in relative abundance sharply with depth. *Acidobacteria*, *Bacteroidetes*, *Latescibacteria*, *Nitrospirae*, *Deferribacteres* and *Gemmatimonadetes* all decreased with depth and accounted for less than 1% of total reads per sample. *Proteobacteria* declined steadily throughout the core from ~25% of total reads initially to ~4% by the end of the core (Fig 2). *Chloroflexi*, *Aminicenantes* and *Planctomycetes* all declined gradually throughout the core profile.

These downcore results suggest that ANME-1 performs either AOM or methanogenesis, but direct evidence for this is best obtained from experimental manipulation of an enrichment culture. Fortunately, ANME-1 enrichments have previously been shown to reverse between net AOM and net methanogenesis based on hydrogen and sulfate availability (2, 25, 34). One ANME-1 enrichment was shown to contain ANME-1 as the sole archaea, as well as SEEP-SRB1 sulfate reducers, and other bacteria (2), and demonstrated that “methane consumption was reversibly inhibited” by hydrogen additions (25). Methane concentrations increased over a period of three days when hydrogen concentrations were high (> 0.5 mM), even when sulfate was present. We calculated the rate of this methane increase and found it to be equivalent to the rate of methane decrease after hydrogen was consumed (Fig. 5). As in most AOM enrichment cultures, AOM was assumed to be driven by the dominant archaea present (ANME-1); we tested whether ANME-1 were also the dominant archaea present when the enrichments switched to net methane production.

**Figure 5.**
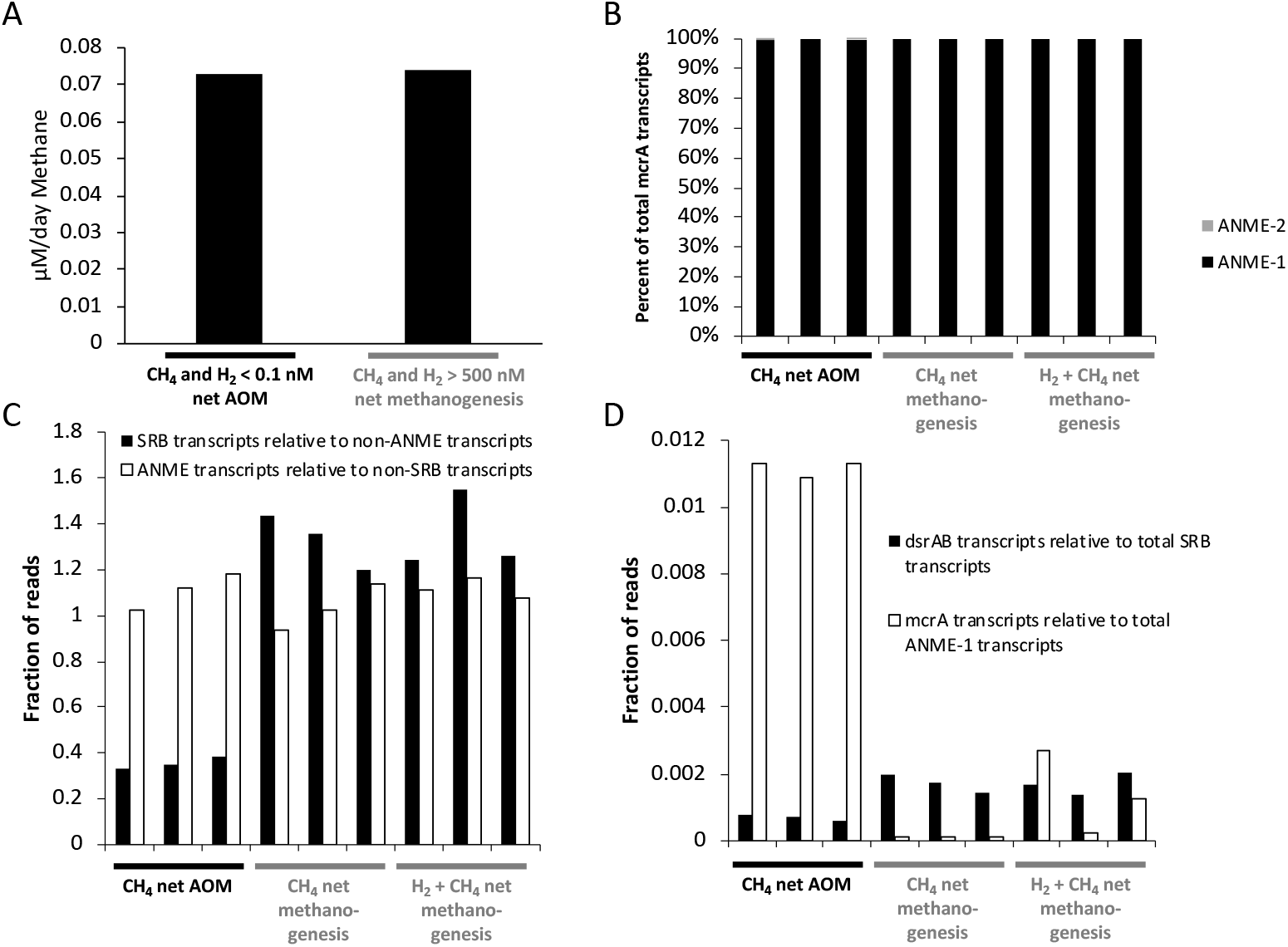
In laboratory enrichments, ANME-1 were likely responsible for AOM in methane-consuming conditions and methanogenesis in methane- producing conditions, controlled by the hydrogen concentration. Shown are A) rates of methane consumption or production, dependent on hydrogen concentrations, B-D) transcript read abundances when triplicates were gassed with methane alone (AOM), hydrogen alone (methanogenic), or hydrogen plus methane (methanogenic), with B) taxonomic identities of all methyl coenzyme-M reductase subunit A (*mcrA)* gene transcripts, C) relative 16S rRNA gene transcript abundance of putative sulfate reducers relative to all transcripts besides those of ANME-1 and of ANME-1 relative to all transcripts besides those of sulfate reducers, and D) transcripts of dissimilatory sulfite reductase subunits A and B (*dsrAB*) relative to total transcripts from sulfate reducing bacteria and transcripts of *mcrA* relative to total transcripts from ANME-1. Data were from A) Wegener et al., 2015 Figure S4, B) our Blast analysis of transcripts obtained from the NCBI short read archive from Wegener et al., 2015, and C and D) Wegener et al., 2015.

We performed a Blast search of *mcrA* genes from all cultured methanogens and uncultured groups containing *mcrA* against the transcriptomes from these enrichments. RNA transcriptomes provide an accounting of all the genes that are being actively transcribed to RNA at the time of sampling, representing abundance and activity. We found that only ANME-1 had hits at an e-value cutoff of <1 × 10^−10^, during both net AOM and net methanogenesis, with the exception of <0.01% hits to ANME-2, another clade of AOM-performing organisms (48) in two of the AOM-performing incubations (Fig. 5). Therefore, the only organism with the genes capable of methane metabolism in each of these incubations was ANME-1, and possibly ANME-2. Wegener et al. suggested that ANME-1 decreased in metabolic activity under conditions of net methanogenesis, since sulfate reducers prefer electrons from hydrogen than from AOM (49). In partial agreement with this interpretation, sulfate-reducing bacteria transcripts and those of dissimilatory sulfite reductase subunits A and B (*dsrAB*) increased relative to non-ANME organisms after hydrogen addition, suggesting sulfate-reducing bacteria were stimulated by the hydrogen additions (Fig. 5c). However, total ANME-1 transcripts did not decrease relative to background populations of other organisms after hydrogen addition (Fig. 5c), suggesting that ANME-1 populations were similarly abundant and active under both AOM and methanogenesis. ANME-1 *mcrA* transcripts, however, were greatly elevated under AOM conditions with low H_2_ (Fig. 5). This is consistent with observations for cultured methanogens, which have been shown to up-regulate *mcrA* gene expression in response to low H_2_ concentration (50, 51). Collectively, these transcript data support active methane metabolism by ANME-1, and only ANME-1, during net AOM and net methanogenesis in these enrichment experiments.

Enrichments of from Hydrate Ridge and Amon Mud Volcano show similar H_2_-dependent reversibility (35). In this study, washing the ANME-1 enrichments free of sulfate was sufficient to see steady methane increases over a period of 30 days, presumably fueled by hydrogen produced by fermentation of organic matter present in the enrichments. The fact that methane started being produced in less than a day and did not increase in production rate over 30 days suggests that the methane was produced by the ANME-1 that were already there, not a growing subpopulation of a different organism, since this would have caused an exponential methane increase and a substantial lag time. Adding H_2_ to the headspace increased the rate 100-fold, further supporting hydrogenotrophic methanogenesis in ANME-1. The rate of methanogenesis in these experiments was much lower than that of AOM, likely because some ANME-1 cells were washed out of the system along with the sulfate. Collectively, these studies demonstrate that the evidence for ANME-1 enrichments performing methanogenesis is as strong as the commonly-accepted evidence that ANME-1 enrichments perform AOM.

If ANME-1 cells perform AOM through a reversible interspecific hydrogen transfer to sulfate-reducing bacteria, then ANME-1 must contain hydrogenases, which are enzymes capable of metabolizing hydrogen. Although ANME-1 genomes contain homologs for the hydrogenases of cultured methanogens, they have thus far been found to lack the active site (23, 25, 49). One possible explanation is that the active subunit of a typical methanogenic hydrogenase is present in ANME-1 genomes, but has not yet been sequenced because the genomes are incomplete, in a similar situation to the *mer* gene which was discovered when more genomes became available (20). Another possibility is that ANME-1 genomes contain a novel hydrogenase active site. All known methanogenic hydrogenases are from cultures grown with extremely high hydrogen concentrations (usually 80% by volume of the headspace). Obligate hydrogenotrophic methanogens undergo major cellular rearrangements due to hydrogen limitation, such as *Methanocaldococcus jannaschii*, which produces flagella when hydrogen is low (17) and alters its central metabolic pathway by decreasing expression of H_2_-dependent methylene-tetrahydromethanoptern (H_4_MPT) dehydrogenase (Hmd), and increasing expression of coenzyme F_420_-dependent methylene-H_4_MPT dehydrogenase (Mtd) (52, 53). This may be because low extracellular hydrogen concentrations create a gradient that causes hydrogen, which is membrane-permeable, to “leak” out of the cell (54), making F_420_ a more efficient electron carrier than H_2_. Adaptation of ANME-1 to low hydrogen conditions may explain the abundance of enzymes utilizing F_420_ rather than hydrogen (23). ANME-1 contains Mtd, which alone is sufficient for metabolizing hydrogen (55). However, the lack of the alpha catalytic subunit suggests that ANME-1 may have a variation of Mtd with high hydrogen affinity, similar to the Hmd_II_ variant of Hmd which has a high hydrogen affinity (56). Accordingly, Beulig et al. 2018 hypothesized that one of the F_420_-reducing hydrogenases could access electrons from hydrogen either alone or in combinations with an adjacent heterodisulfide reductase or a Nuo-like oxidoreductase complex (20).

If ANME-1 has a reversible metabolism, then all the enzymes in the pathway must be reversible. Cells often ensure that metabolic reactions that are essential to the cell, yet operate at ΔG close to zero, only flow forward by employing “irreversible” enzymes. Such enzymes, such as phosphofructokinase in glycolysis, bind their product at an allosteric site away from the active site in order to disable the enzyme activity when product builds up, in order to prevent back-flow. However, all the genes of methanogenesis have been shown to be reversible (57); possibly because they are well-poised to catalyze whichever direction is exergonic. The high degree of reversibility of the hydrogenotrophic methanogenic enzymatic pathway is shown by the fact that hydrogenotrophic methanogens express isotopic fractionation factors highly dependent on the free energy available for the reaction (52). Therefore, ANME-1 may gain energy through AOM in the SMTZ, increasing their cell abundance relative to other methanogens. Then, when hydrogen concentrations increase after sulfate is depleted, ANME-1 have a head start on these other methanogens, and can outcompete them for meager resources.

We conclude that ANME-1 performs methanogenesis in the MZ, because it is the dominant organism with the genetic capability to do so in both geochemically-characterized White Oak River estuary sediments and well-controlled laboratory experiments. This either occurs through a metabolic reversal depending on sulfate-reducing bacterial control of hydrogen concentrations (Fig. 6), or it is caused by subclades with identical 16S rRNA sequences that drive either AOM or methanogenesis. However, it does not appear to occur through a major die-off of the majority ANME-1 population at the base of the SMTZ. Our conclusion is further supported by geochemical models (12, 13), other ANME-1 enrichments (25, 35), reversibility of the methanogenic biochemical pathway (20), and dominance of ANME-1 in common marine sediments in geographically widespread areas (8, 19, 20). Although known methanogenic clades have been cultured from marine sediments and identified by gene surveys targeting cultured clades (58), we are not aware of any study that identified cultured methanogens in methanogenic marine sediments when using universal primers, except for salt marsh tidal flats which are periodically exposed to air (59). However, many studies of deep-sea sediments lack both ANME-1 and cultured methanogens (60, 61), meaning that our results should not be extrapolated to all deep-sea locations. Marine sediments are the third largest producers of methane on Earth, after rice production and wetlands, but they are only the ninth largest methane emitters (1). If ANME-1 archaea are responsible for methanogenesis in many marine sediments, then they are some of the dominant methane emitters on Earth, in addition their better-known function as some of Earth’s most efficient methane oxidizers. The prevalence of metabolically reversible ANME-1 on Earth suggests that this level of extreme metabolic flexibility may be a more widespread feature of organisms specialized to survive in ultra-low energy environments. This could be used as a guide in the search for habitable places on Earth and extraterrestrial environments.

**Figure 6.**
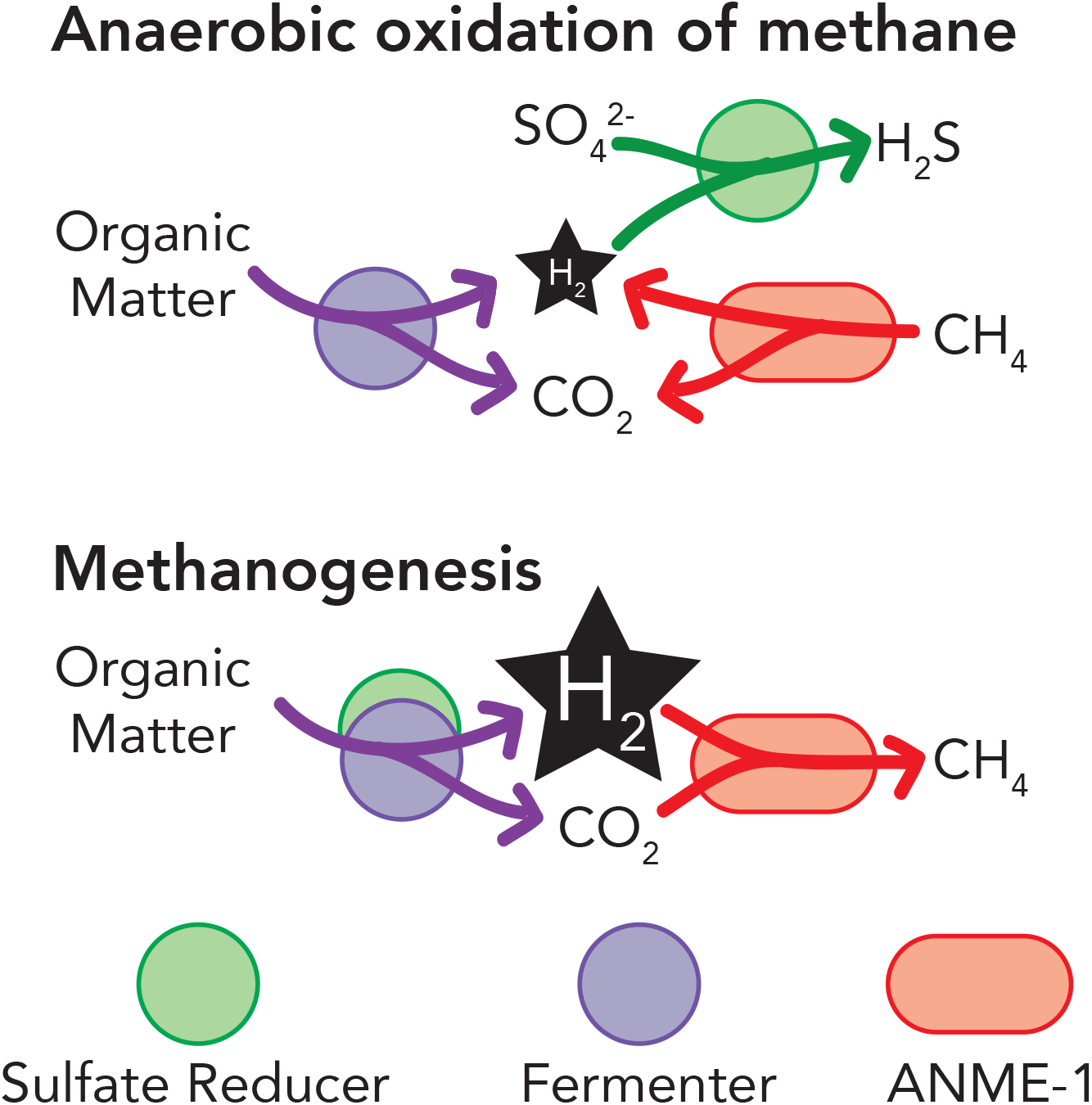
Model for AOM and methanogenesis in non-seep marine sediments. Fermenters produce hydrogen from organic matter under both conditions. During anaerobic oxidation of methane, sulfate reducers bring hydrogen concentrations so low that methane oxidation is exergonic. During methanogenesis, sulfate reducers have run out of sulfate so hydrogen concentrations build up to a level that makes ANME-1 switch to forward methanogenesis. Sulfate reducers may persist by switching to fermentation. Purple = fermenters, green = sulfate reducers, and red = ANME-1.

## Acknowledgments

We thank Marc Alperin for invaluable conversations about methane over 17 years, Brett Baker for conversations about the metagenomic data, Andrew Steen, Jordan Bird, Lauren Mullen, Taylor Royalty, Katherine Sipes, Katherine Fullerton, and Joy Buongiorno for assistance retrieving the sediment samples, Michael Piehler for the use of his laboratory at the UNC Institute of Marine Sciences, Frank Löffler for use of ion and flame ionized detector chromatographs, Robert Murdoch and the University of Tennessee Knoxville Bioinformatics Resource Center for computational resources, and Christopher G. Kevorkian and Alexandra Emmons for aid in analysis. This work was funded by Alfred P. Sloan Foundation Fellowship (FG-2015-65399) and NASA Exobiology (NNX16AL59G).

## Methods

### Sample collection and geochemistry

Two push cores (core 1 was 78 cm deep and core 6 was 75 cm deep) were collected in May 2017 (1 m) at the White Oak River Estuary Station H (34 44.490’ N, 77 07.44’ W), in 1.5 m water depth. Other cores (accounting for numbers 2-5) were taken and used for other projects or thrown away because they were not long enough. Using a plunger inserted from the bottom to push the core up, sediment intervals were sectioned in 3 cm intervals from 0 to 9 cm, 1 cm intervals to 60 cm and in 3 cm intervals below that. These intervals were chosen to obtain high depth resolution in the section most likely to include anaerobic oxidation of methane. Methane, sulfate, δ^13^C of methane, hydrogen, and cell counts were measured as described previously (29). To measure methane, sediment was quickly sub-cored with a plastic cut-off 4 ml syringe, placed into glass serum vial containing 1 ml 0.1 M KOH, sealed with a butyl rubber stopper, and shaken to mix. Methane was determined by injecting 100 μl of gas from the headspace, after shaking the bottle vigorously for 1 min, into a gas chromatograph with a flame ionization detector (Agilent, Santa Clara, CA). Due to volume limitations when slicing into 1 cm intervals, porosity was not measured in these cores, but was used from core A in a previous core (8). To measure δ^13^C values of methane, 4 ml of headspace from the vial used for methane measurements was removed via syringe and injected into a gas bag containing hydrocarbon free zero gas (Airgas, Radnor, PA). This was then measured on a cavity ring down spectrometer using a small sample introduction module (Picarro, Santa Clara, CA). To measure hydrogen, 2 ml of carefully sub-cored sediment was placed into a 10 ml brown glass vial being careful to disturb the sediment as little as possible, sealed with a black butyl stopper, and gassed with helium. The negative control was an empty bottle gassed with helium. After 4 days equilibration at near *in situ* temperature (21°C), 500μl of headspace gas was injected into a Peak Performer 1 Reducing Compound Photometer (Peak Laboratories, Mountain View, CA). Premixed hydrogen ppm lab bottles (Airgas) were used as standards. For cell abundance, 1 ml of sediment was placed in a 2 ml screw cap tube with 3% paraformaldehyde. To measure sulfate, a 15 ml plastic tube was filled completely with sediment and centrifuged at 5,000 ×g for 5 minutes. A syringe was used to remove the supernatant below the air interface. The porewater was filtered using a 0.2 μm syringe filter into 100 μl of 10% HCl to a final volume of 1 ml. Porewater sulfate concentrations were determined by ion chromatography (Dionex, Sunnyvale, CA). A 10 ml cut off syringe was filled, capped with parafilm, and frozen at −80°C for later molecular analysis.

### Cell quantification

Total cell counts were determined by direct epifluorescence microscopy using SYBRGold DNA stain (Invitrogen, Carlsbad, CA). Sediments were sonicated at 20% power for 40 seconds to disaggregate cells from sediments and diluted 40-fold into PBS prior to filtration onto a 0.2 μm polycarbonate filter (Fisher Scientific, Waltham, MA) and mounted onto a slide.

### 16S Ribosomal RNA Gene Amplicons

DNA was extracted from frozen sediments using the Fast DNA kit for Soil (MP Bio, Santa Ana, CA). The V4 region was amplified using primers 806r and 515f (62), as a universal primer pair for Bacteria and Archaea. A negative sample containing DNA extracted from autoclaved sediment control yielded no amplification. Library preparations via Nexterra kit and sequencing using an Illumina MiSeq were performed at the Center for Environmental Biotechnology at the University of Tennessee in Knoxville, producing 14,162,094 reads total. The CLC Genomic Workbench 10.0 (CLC Bio, a QIAGEN Company, Aarhus, Denmark) was used to trim adaptors and make contigs of bidirectional sequences, cluster operational taxonomic units (OTUs) at 97% similarity, and classify them with the Silva reference set 132 (63). 36.4% of a total of 866,834 unique sequences were removed as chimeric, resulting in a total of 9,165,958 reads in 25,116 OTUs. Approximately 5% of the remaining sequences were removed for not classifying as Archaea or Bacteria. Reads were then randomly subset from sample libraries to the size of the smallest library (77,609) for normalization. Only OTUs with at least an average of 2 reads per sample were considered, resulting in 2,307 OTUs across 59 libraries. 16S rRNA genes from metagenomes from this site (37) were obtained from IMG/ER, and classified analogously to the amplicon dataset.

### Data analysis of published ANME-1 enrichment experiments

All nine transcriptomes from Wegener et al., 2015, were downloaded from the NCBI SRA as fastq files, and trimmed for quality with Trimmomatic 0.33 (64). Twenty-six genes for methyl coenzyme M reductase were obtained from NCBI to make a custom Blast database including representatives from all major cultured methanogens, and mcrA genes from MAGs of uncultured organisms. The mcrA database included the following mcrA genes: *Methanospirillum hungatei* strain JF-1 (AF313805), *Methanobacterium bryantii* strain DSM 863 (AF313806), Uncultured Methanobacteriaceae methanogen RS-MCR12 (AF313818), Uncultured RC1 methanogen RS-MCR15 (AF313821), *Methanoculleus bourgensis* strain DSM 3045 (AF414036), *Methanosaeta concilii* strain DSM 3671 (AF414037), *Methanocaldococcus jannaschii* strain DSM 2661 (AF414040), *Methanopyrus kandleri* strain DSM 6324 (AF414042), *Methanosphaera stadtmanae* strain DSM 3091 (AF414047), *Methanothermococcus thermolithotrophicus* strain DSM 2095 (AF414048), *Methanocorpusculum labreanum* (AY260441), Uncultured ANME-1Guaymas archaeon (FR682814), Uncultured ANME-1 archaeon clone F17.1_30A02 (AY324363), Uncultured ANME-1 archaeon clone GZfos_17_30.54 (AY324369), Uncultured ANME-2 archaeon clone GZfos_26_28.10 (AY324370), Uncultured ANME-2 archaeon clone GZfos_35_28.12 (AY324371), *Methanomassiliicoccus luminyensis* strain B10 (HQ896500), Uncultured Methanomicrobiales archaeon clone H07 (AY837764), Uncultured Methanomicrobiales archaeon clone B12 (AY837766), Uncultured Methanomicrobiales archaeon clone C01 (AY837767), Uncultured Methanosarcinales archaeon clone C05 (AY837769), Uncultured ANME-2Guaymas archaeon clone D06 (AY837771), Uncultured Methanosarcinales archaeon clone B09 (AY837774), Uncultured Methanococcales archaeon clone D03 (AY837775), *Methanosarcina mazeii* strain C 16 (KT387805).

The NCBI Blast tool was used to obtain hits with e-values less than 1e-10 of the transcriptomes among this *mcrA* database. Rates of methane production and consumption were calculated through WebPlotDigitizer on Extended Data Figure 4 from Wegener et al., 2015, which showed methane concentrations through time when the added hydrogen concentrations were high (> 0.5 mM) or low (< 0.1 mM).

### Data archiving

16S rRNA gene sequences can be found at the NCBI Genbank short read archive with accession number PRJNA565996.

